# Structure-based Antibody Renumbering

**DOI:** 10.64898/2026.09.16.751788

**Authors:** Diego del Alamo

## Abstract

Antibody numbering schemes like IMGT and Chothia assign each residue in the variable domain a consistent index based on substructural position. These annotations standardize sequences with different lengths, facilitating tasks ranging from engineering of individual molecules during drug development to large-scale curation of training data for *de novo* antibody design. Yet existing algorithms for performing this annotation process, termed renumbering, rely exclusively on amino acid sequence for inference. Consequently, these can fail when presented with unnatural or unusual features such as long CDRs or engineered insertions. To address this gap, this work introduces Structure-based Antibody Renumbering, abbreviated SAbR, a method that assigns these annotations from structure alone. SAbR shows comparable performance to sequence-based renumbering methods on held-out expert-annotated structures, as well as high agreement with sequence-based methods in a larger benchmark of diverse structures. It also outperforms peer methods on *de novo*-designed molecules with long CDR loops, and shows higher success rates than renumbering by structural alignment. However, limited generalization performance is observed in more distantly related systems. Overall, these results establish structure-based renumbering as a robust alternative for natural and engineered antibodies when such data is available. Code and model weights are available on [GitHub](https://github.com/delalamo/SAbR).

## 1 Introduction

Variable domains of antibodies consist of both a conserved immunoglobulin fold as well as loops with diverse lengths and conformations [1]. Numbering schemes such as IMGT, Chothia, Kabat and others assign each residue an index based on its position in this fold [2–10]. This facilitates consistent comparisons across molecules of different lengths. Although these schemes are structurally motivated, current renumbering tools operate exclusively on sequence [11, 12]. For example, they might annotate residues in CDRs by detecting conserved residues at their N- and C-termini, rather than the geometry of those residues within the overall domain. As such, these tools can fail when, for example, CDRs are unusually long or contain unnatural sequence features that confound such detection schemes.

In this communication, I introduce Structure-based Antibody Renumbering, or SAbR for short, a tool for inferring residue assignments from backbone heavy-atom coordinates without using sequence (Figure 1). SAbR reproduces annotations from expert-annotated antibody structures, and shows high agreement with sequence-based methods in a larger benchmark, including chains with long CDRs. It also outperforms sequence-based baseline methods on *de novo*-designed antibodies with long CDR loops. Although parametrized only on predicted mammalian antibody structures, it shows some utility in renumbering the framework regions of distantly related systems with low sequence homology, although broad generalization remained limited and contingent on template selection. Finally, SAbR outperforms structure-based alignment methods like TM-align that can be repurposed for annotation transfer. These results establish the viability of structure-based renumbering when such data are available.

**Figure 1:**
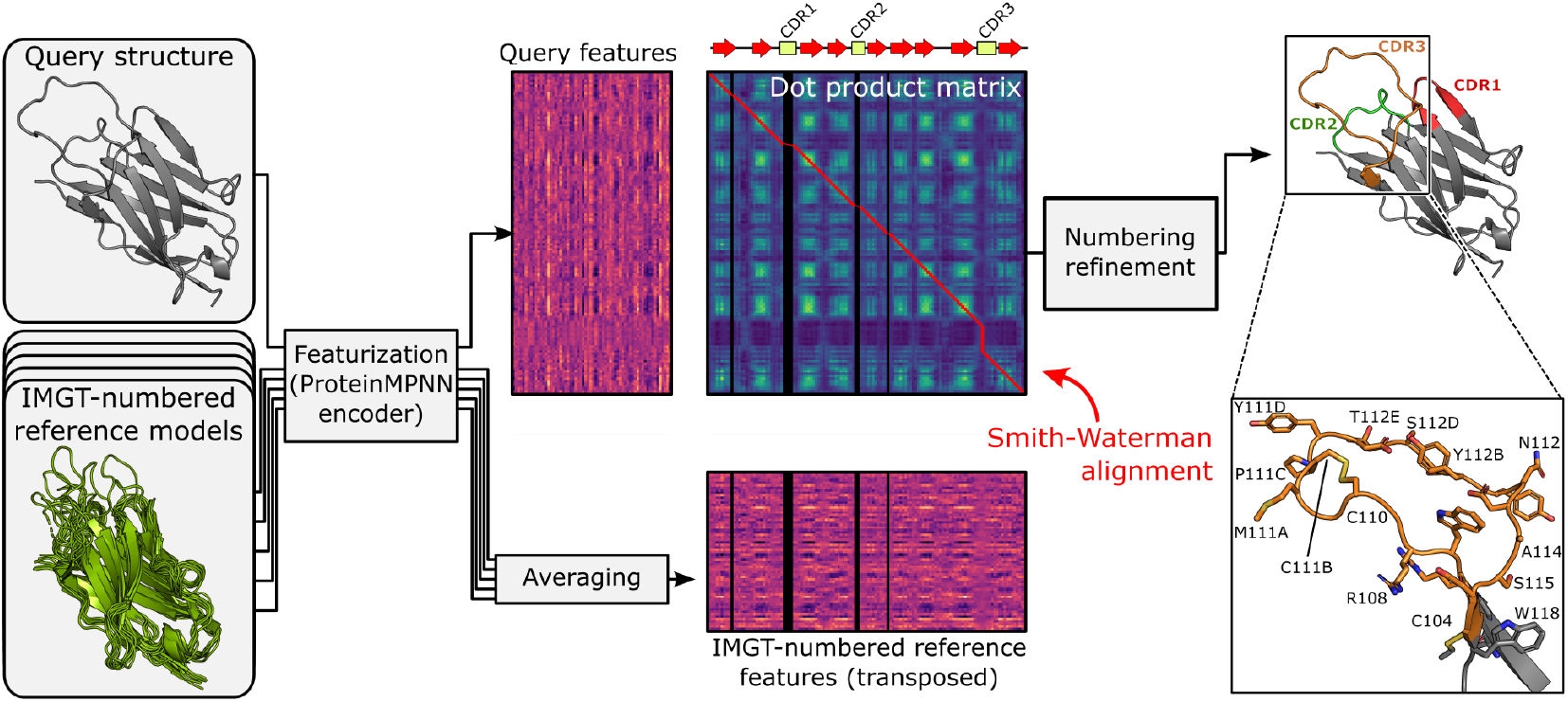
SAbR renumbers antibody structures by comparing their per-residue ProteinMPNN embeddings to average embeddings calculated from thousands of reference models.

## 2 Related work

### Antibody variable domains and numbering schemes

Antibody variable domains adopt a beta-sandwich immunoglobulin fold consisting of nine beta strands comprising the framework and three variable-length CDR loops [13]. Diverse sequence lengths complicate mapping of structural features to consistent residue indices. Like other protein families [14–16], antibodies have specialized numbering schemes that attempt to map structural features to standardized indices. A recent review [1] discusses the origins and relative merits of these schemes.

### Sequence-based renumbering

Current sequence-based renumbering methods mostly repurpose algorithms used for sequence alignment. RIOT aligns the V- and J-gene segments of queries directly to numbered germline references [17], while AbNum and AntPack use position-specific profiles of framework sequence conservation [9, 18] and ANARCI uses hidden Markov models [11, 19]. A notable exception is ANARCII, which eschews alignment entirely and renumbers using transformer neural networks trained on ANARCI-numbered antibody sequences [12]. These tools are instrumental in curation of renumbered antibody structures in databases like SAbDab [20].

### Structure-based alignment

Structure-based methods align residues by backbone geometry rather than sequence. TM-align combines secondary-structure assignment and superposition and serves as the alignment engine for immunoglobulin-based numbering annotation approach IgStrand [21, 22]. SoftAlign instead aligns residue representations from a retrained version of the graph neural network ProteinMPNN, achieving performance competitive with TM-align [23, 24]. Foldseek uses tokenized representations of residues for fast structure search [25].

## 3 Methods

### 3.1 Problem formulation and method overview

SAbR builds on SoftAlign, which uses a ProteinMPNN encoder to generate residue representations for queries that are aligned to those of reference templates with differentiable Smith–Waterman alignment [23, 24, 26]. As with the sequence-based antibody renumbering tools outlined above, numbering annotations are transferred through this alignment. This offers several avenues for improving renumbering performance by leveraging large numbers of templates. Each template may be aligned to the query independently, with best-of-k selection returning the one with the highest score (Figure 2A). Alternatively, individual residue assignments from independent alignments can be jointly considered using a consensus or majority-vote approach, which was found to be more performant (Figure 2B). Finally, the ProteinMPNN representations of consistently numbered reference structures can be averaged position-wise. This yields exactly the mean of their reference-specific dot-product score matrices while requiring only one alignment calculation. Figure 2C shows that this strategy recovers most of the performance gains from ensembling independent alignments while scaling better to more templates.

**Figure 2:**
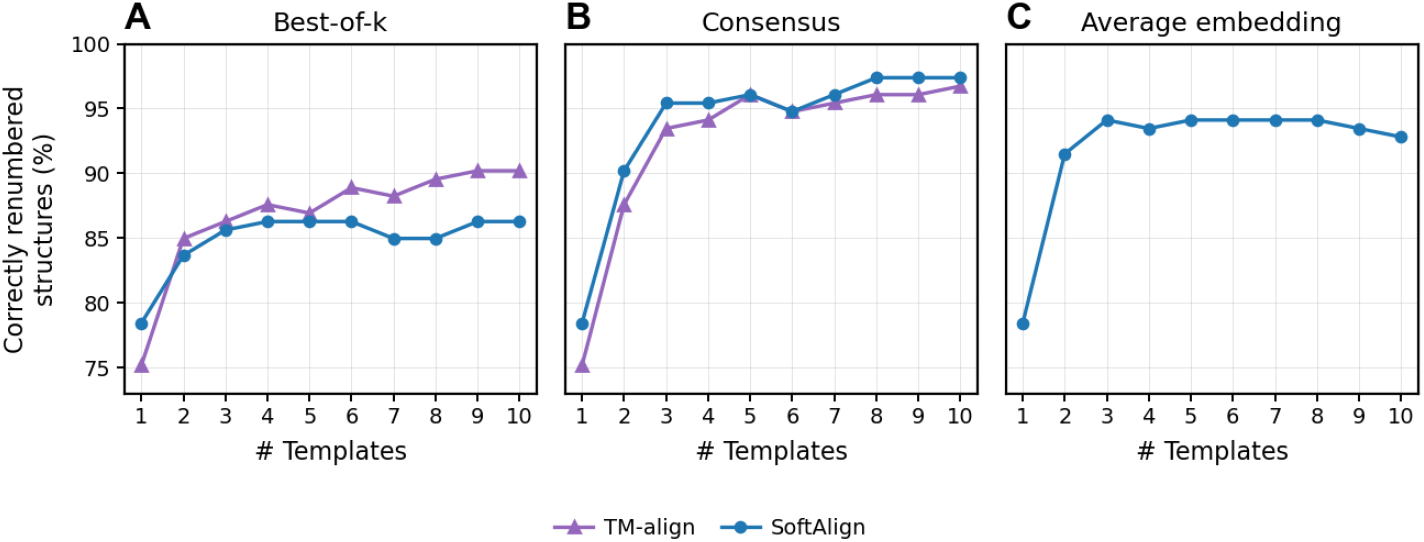
Multi-template alignments improve renumbering performance. Alignment accuracy shown for 153 SAbDab test structures with ≥ 10 homologs in the pre-cutoff dataset. Templates were retrieved based on sequence identity to the query. TM-align was omitted from panel C as it does not use embeddings for alignment.

### 3.2 IMGT-numbered reference embeddings

Construction of separate averaged embeddings from reference structures for heavy, kappa, and lambda chains requires that each IMGT index corresponds to a consistent structural position across all structures. Rather than relying on deposited experimentally determined structures, SAbR makes use of over 100,000 publicly available IMGT-numbered ImmuneBuilder models of paired antibodies consisting of 148,832 heavy chains, 113,269 kappa chains, and 35,541 lambda chains [27, 28]. Figure 3 shows how framework residues in these models, more so than those in deposited experimental structures, aligned almost exclusively with residues carrying the same IMGT index.

**Figure 3:**
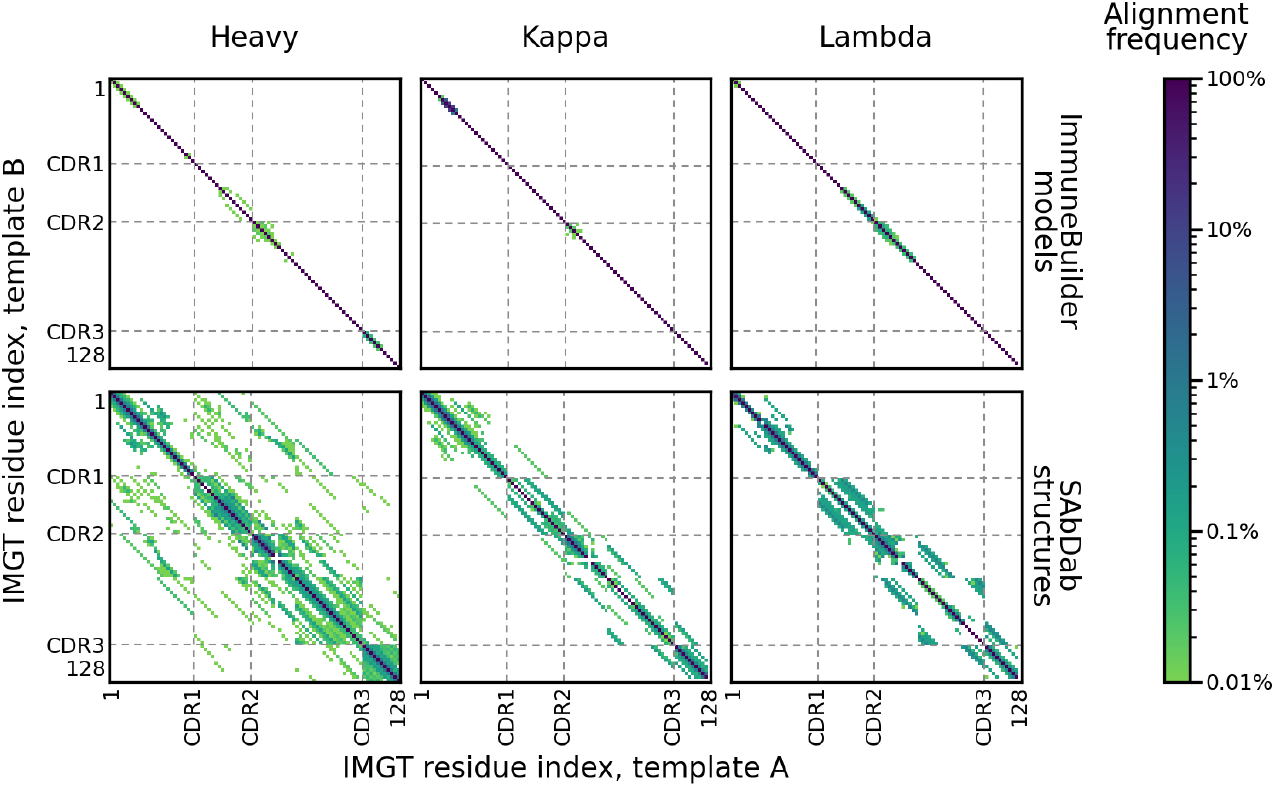
Structural alignments of ImmuneBuilder models, but not SAbDab structures, are concentrated along matching IMGT indices. Symmetrized heatmaps show alignment frequencies between 10,000 representative pairs of IMGT-renumbered heavy, kappa, and lambda-chain structures from both datasets. CDRs and residues with insertion codes are not shown. ImmuneBuilder models show fewer off-diagonal matches, suggesting greater adherence to the structural motivation of the IMGT numbering scheme.

When averaging the ProteinMPNN representations of these ImmuneBuilder templates, residues were excluded if they either had insertion codes present, or were found in fewer than 30% of structures (Figure S1).

### 3.3 Gap penalties and fine-tuning

The differentiable Smith–Waterman alignment algorithm used by SAbR and SoftAlign relies on gap open penalty *γ*_o_ and gap extension penalty *γ*_e_, which are jointly learned with the weights of its ProteinMPNN encoder [23]. Surprisingly, its released parameters include a positive gap extension penalty (i.e., a bonus), which rewards the extension of an existing gap and sometimes spuriously introduced massive insertions into multidomain structures (example shown in Figure S2). Therefore, in addition to evaluating the base SoftAlign weights, this study evaluates a fine-tuned version of SoftAlign with non-positive gap penalty values. Fine-tuning of both the learned gap penalty values and the accompanying ProteinMPNN encoder proceeded for one epoch using a learning rate of 3 *×* 10^−5^ and the SoftAlign training set, and the non-positivity constraint on *γ*_e_ was introduced over *T* = 500 steps. At training step *t*, the effective gap-extension score was set to

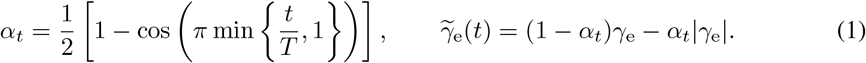

Thus, the original value is used at the beginning of fine-tuning, while 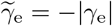 once the constraint is fully applied. The gap open penalty was similarly constrained, but was already negative in the base weights. The same optimizer settings as the original training run were reused (Adam optimizer, *β*_1_ = 0.9, *β*_2_ = 0.999).

### 3.4 Numbering reconstruction and post-processing

Alignments are run with a temperature of 10^−4^. SAbR assigns the query to the chain type with the highest alignment score and selects the corresponding alignment for numbering. The selected alignment was discretized and mapped onto the full 128-position IMGT reference. Residue assignments within the CDRs were then reconstructed deterministically following IMGT convention using prespecified residue pairs as framework anchors: IMGT positions 23/40 for CDR1, 54/67 for CDR2, and 104/118 for CDR3 [7]. These anchors deviate from those used by the IMGT numbering scheme by default, and were selected on the basis of conservation of structure rather than sequence. Figure 4 shows how this deterministic correction maintains accurate assignments as CDR length increases.

**Figure 4:**
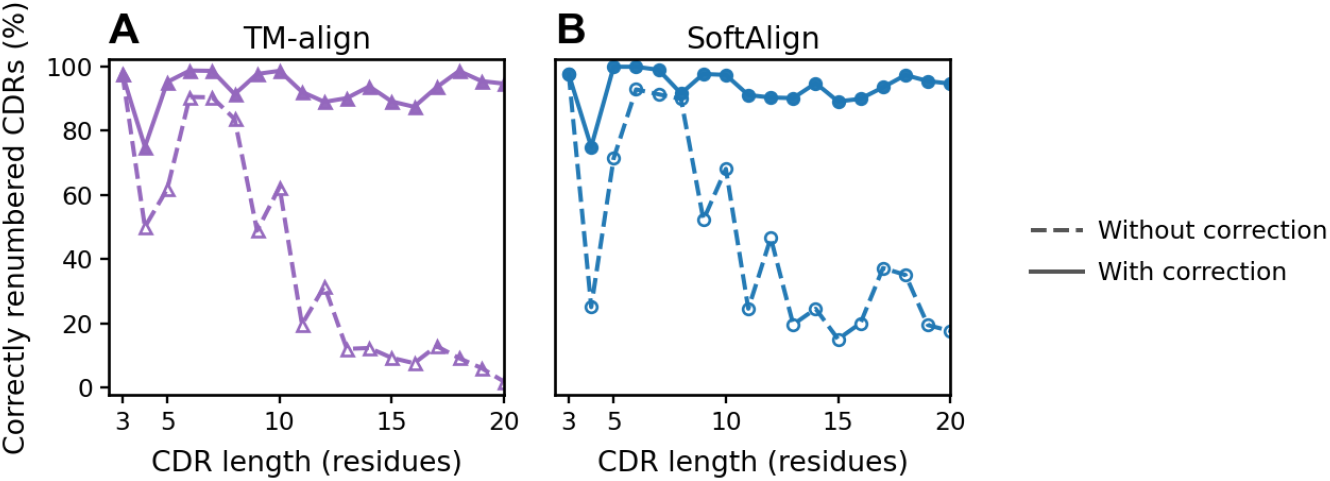
Renumbering by annotation transfer alone is error-prone for long CDRs. CDR-specific renumbering errors can be avoided with deterministic renumbering following IMGT conventions. Results obtained with 1,141 SAbDab test set structures.

The DE loop was similarly reconstructed between positions 79 and 85, with insertions assigned to index 82. The corrected alignment was converted into an ANARCI-compatible sequence of match, deletion, and insertion states [11, 19]. Unoccupied reference positions became deletion states, whereas query residues located between aligned positions became insertion states. IMGT insertion-code conventions are applied, and the resulting assignments are then translated into Chothia, Kabat, Martin, AHo, or Wolfguy numbering [2, 4, 5, 8–10].

### 3.5 Renumbering multidomain structures

SAbR takes entire chains as input, including leader peptides and constant domains. To renumber chains with multiple variable domains, such as scFvs, reference embeddings were concatenated, and to account for the presence of the linker, gap penalties between residues aligned to the terminal edges of these domains were set to zero. This permitted the alignment to freely insert gaps in the linker region as needed without relying on user-specified domain boundaries. This scheme worked for other multi-domain chain arrangements as well.

## 4 Experiment details

### Baseline methods

Sequence-based antibody numbering methods included ANARCI and ANARCII [11, 12]. Structure-based methods included TM-align, SoftAlign, and Foldseek [21, 23, 25]. The first two used templates identified by sequence identity; MMseqs2 was used to identify the pre-cutoff template with the highest sequence identity to each query, which was then used for structural alignment and annotation transfer [29]. Foldseek used templates identified using either its 3Di alphabet score or 3Di+sequence alignment scores (shown as 3Di+AA) [25].

### Datasets and splits

Two antibody datasets were used for testing: a small-scale, expert-annotated dataset from IMGT/3D-StructureDB [30], and a larger-scale set from SAbDab, whose annotations are largely but not entirely assigned automatically with ANARCI [11, 20]. The former measures renumbering accuracy, while performance on the latter reports renumbering agreement with sequence-based tools. Since ImmuneBuilder was trained on PDB structures deposited through 31 July 2021, this cutoff date was likewise used here [27, 28]. This also post-dates the cutoff for the SCOPe 2.01 database used to train and fine-tune SoftAlign [31]. Post-cutoff structures were restricted to ≤ 3.5 Å resolution and excluded if they were shark antibodies, had missing atoms or residues within the variable domain, or shared ≥ 85% variable-domain sequence identity with any pre-cutoff structure. The remainder of SAbDab structures were clustered at 85% variable-domain sequence identity using MMseqs2 [29], yielding 496 structures with a C-terminal CH1 domain and 645 without. Due to the smaller size of the IMGT/3D-StructureDB dataset, even before clustering, only 42 unique structures were obtained. Analogous filtering and clustering of pre-cutoff structures from SAbDab produced the TM-align and SoftAlign template set.

### Out-of-distribution benchmarks

Five additional systems probed multidomain renumbering, sequence novelty, and structural novelty (Figure 5). The multidomain benchmark comprised 57 post-cutoff experimental scFvs and 666 synthetic scFvs generated from the SAbDab test set with (G_4_S)_3_ linkers modeled using PDBFixer and OpenMM [32, 33]. The sequence novelty benchmark comprised 36 shark VNARs from SAbDab and 946 T-cell receptor (TCR) chains from IMGT/3Dstructure-DB [30], both of which share low sequence identity with conventional antibodies [34–36]. The structural novelty benchmark comprised 37 bovine heavy chains, whose CDRH3 loops can be highly structured and exceed 40 residues in length [37, 38], and 3,380 *de novo* designs with long CDR loops generated from 82 test-set camelid nanobody templates using RFantibody both with and without sequences designed with AbMPNN [39, 40]. Of the latter designs, 2,560 had one CDR redesigned and 820 had all three redesigned; CDR1 and CDR2 lengths ranged from 10 to 50 residues and CDR3 lengths from 11 to 100.

**Figure 5:**
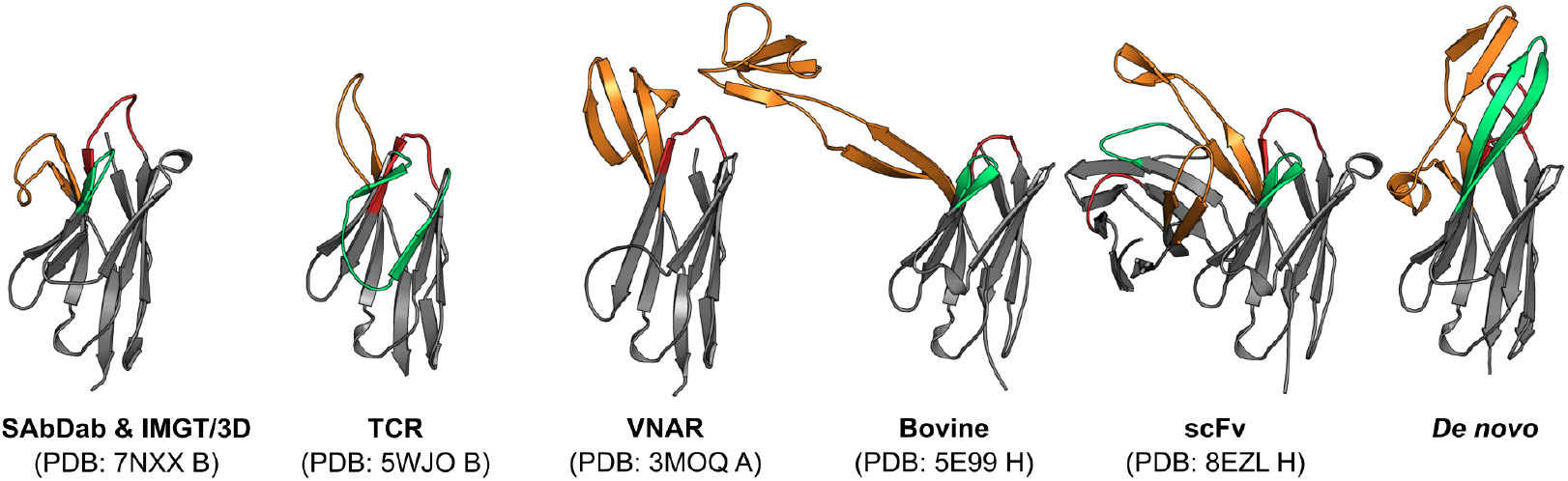
Test case representatives for evaluation of antibody renumbering methods. CDRs 1, 2, and 3 are shown in red, teal, and orange, respectively.

### Evaluation metrics

Successful numbering was primarily evaluated by measuring the number of test set structures that were correctly renumbered in full. Per-residue success was also considered and reported in Table S1. As discussed earlier, all three CDRs and the DE loop (IMGT range 81–84) can be renumbered deterministically from framework anchors, and were therefore excluded from the primary accuracy and agreement metrics. For VNARs, residues 45–75, which constitute a loop replacing CDR2 and its flanking beta strands, were also excluded. For TCRs, the C^*′′*^ strand was omitted due to an off-by-one misalignment with antibody frameworks. IMGT position 128 was also excluded where appropriate. Full-variable-region results including these regions are reported in Table S2.

## 5 Results

### 5.1 Overall performance

Table 1 shows the fraction of structures whose frameworks were correctly renumbered in full by each tested method across each dataset; full variable domain performance is reported in Table S2. SAbR generally matched or outperformed peer methods, with the exception of TCRs, discussed below. As expected, sequence-based methods excelled on generic antibody structures and TCRs, while showing mixed performance on out-of-distribution systems such as *de novo*-designed antibodies. ANARCI for example is not parametrized for VNARs, whereas ANARCII specifically received bespoke training on these sequences [12].

**Table 1:**
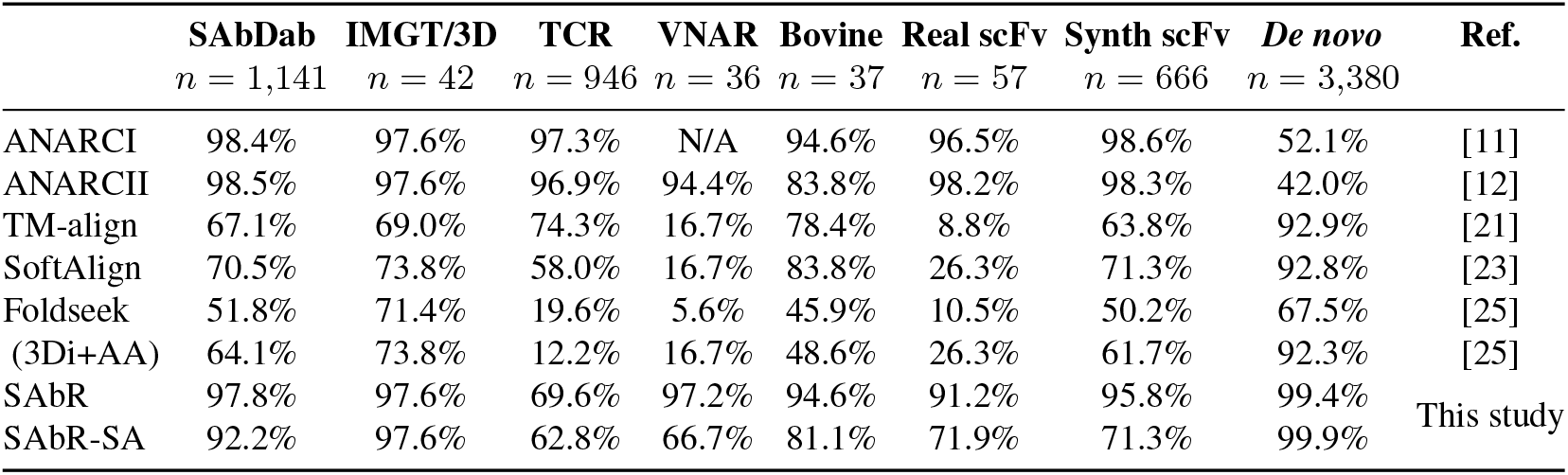
Percentage of test set structures with identical numbering to reference annotations. Single-template results shown for TM-align and SoftAlign. All structure-based TCR and VNAR results use kappa-chain templates. SAbR-SA denotes SAbR with SoftAlign weights. *De novo* results show results only for AbMPNN-designed sequences.

Most structure-based alignment methods, in contrast, excelled on cases where frameworks closely matched the template set, such as bovine and *de novo* designed antibodies with long loops. On *de novo*-designed molecules, structure-based methods demonstrated superior renumbering performance, and were resilient to both CDR lengths and sequence composition (Figure 6, Table S3). Sequence-based methods fared poorly on CDRs that were either unusually long or were not designed (e.g., polyglycine), and showed uneven performance on frameworks with multiple designed CDRs.

**Figure 6:**
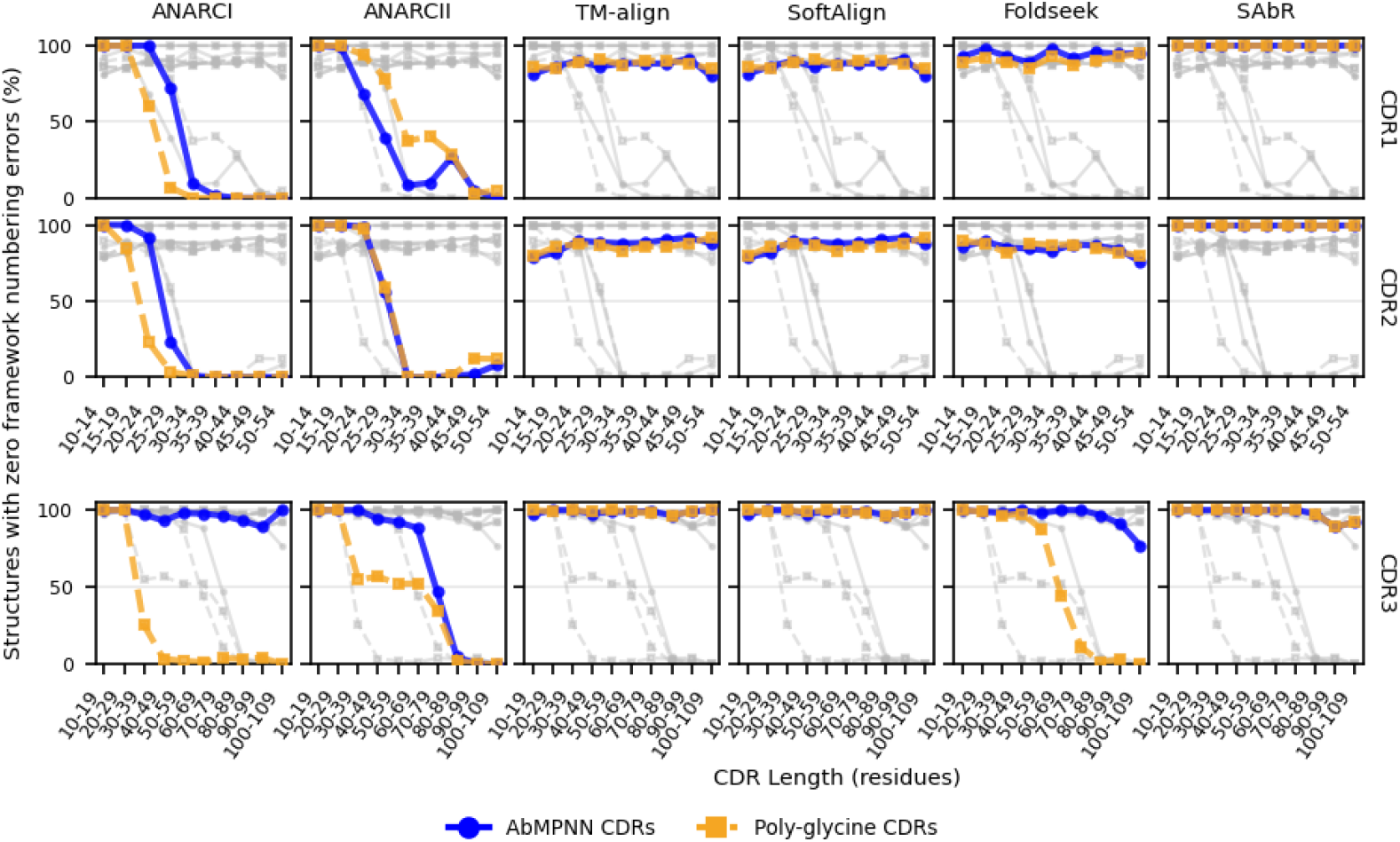
Renumbering performance on synthetic *de novo*-designed antibody structures with only one redesigned CDR. Gray lines indicate other methods for reference. Performance differences between AbMPNN and poly-glycine designs in TM-align and SoftAlign were due to template selection. Foldseek shows 3Di+AA performance.

Structure-based methods additionally showed limited generalization to VNARs but not TCRs. The overwhelming source of error was IMGT position 10, which is absent from heavy and lambda chains and some but not all TCRs [8], and whose existence in the query structure did not appear to reliably influence template selection (Table S4). Overall, these results argue against universal numbering transferability of representations across related but distinct systems.

Beyond this, structure-based alignment methods showed limited performance in scFvs, frequently correctly renumbering only one domain (Table S5), with errors overwhelmingly occurring in the linker.

### 5.2 Ablations and weaknesses

Replacement of default SAbR weights with SoftAlign weights degraded performance in scFvs and test set proteins with the CH1 constant domain present (Table S6), demonstrating how the fine-tuned weights improved renumbering accuracy. However, this fine-tuning did slightly degrade performance on *de novo*-designed test cases.

Two further observations warrant mention. First, compared to sequence-based peer methods, SAbR showed greater sensitivity to the presence of structural gaps, with performance degrading substantially as the number of unresolved residues increased (Figure S3). Second, SAbR’s reliance on alignment scores to auto-detect chain type showed mixed performance, with heavy, kappa, and lambda chains being correctly identified 100.0%, 97.6%, and 88.6% of cases, respectively. Interestingly, all misidentified lambda chains were annotated as heavy chains, but were correctly renumbered anyways. This suggested that chain-specific features, such as the longer DE loop of heavy chains and the bulge at IMGT position 10 of kappa chains, were generally sufficient to delineate these identities, and, in any event, were not strictly necessary for the examined task.

### 5.3 PDB case studies

Beyond the test cases presented here, the PDB contains a veritable zoo of one-off structurally anomalous antibodies against which SAbR was also tested, with some cherry-picked exemplars shown in Figure 7A. PDBs 8SVE and 8RY2 contain ≥ 100-residue cytokine inserts grafted into CDR1, while PDBs 6Y1R and 6Y0E contain different lanthanide binding tag insertions in non-CDR loops. Other examples include long DE loop insertions (PDBs 9BHH and 6NNJ), multi-variable-domain antibody formats (PDBs 7Y6K and 8DT8), and domain-swapped camelid nanobodies (PDB 8V9W), among others. With the exception of the last case, SAbR correctly identified and renumbered insertions, the latter failure being likely due to the disruption to the overall fold caused by dimerization (Figure 7B).

**Figure 7:**
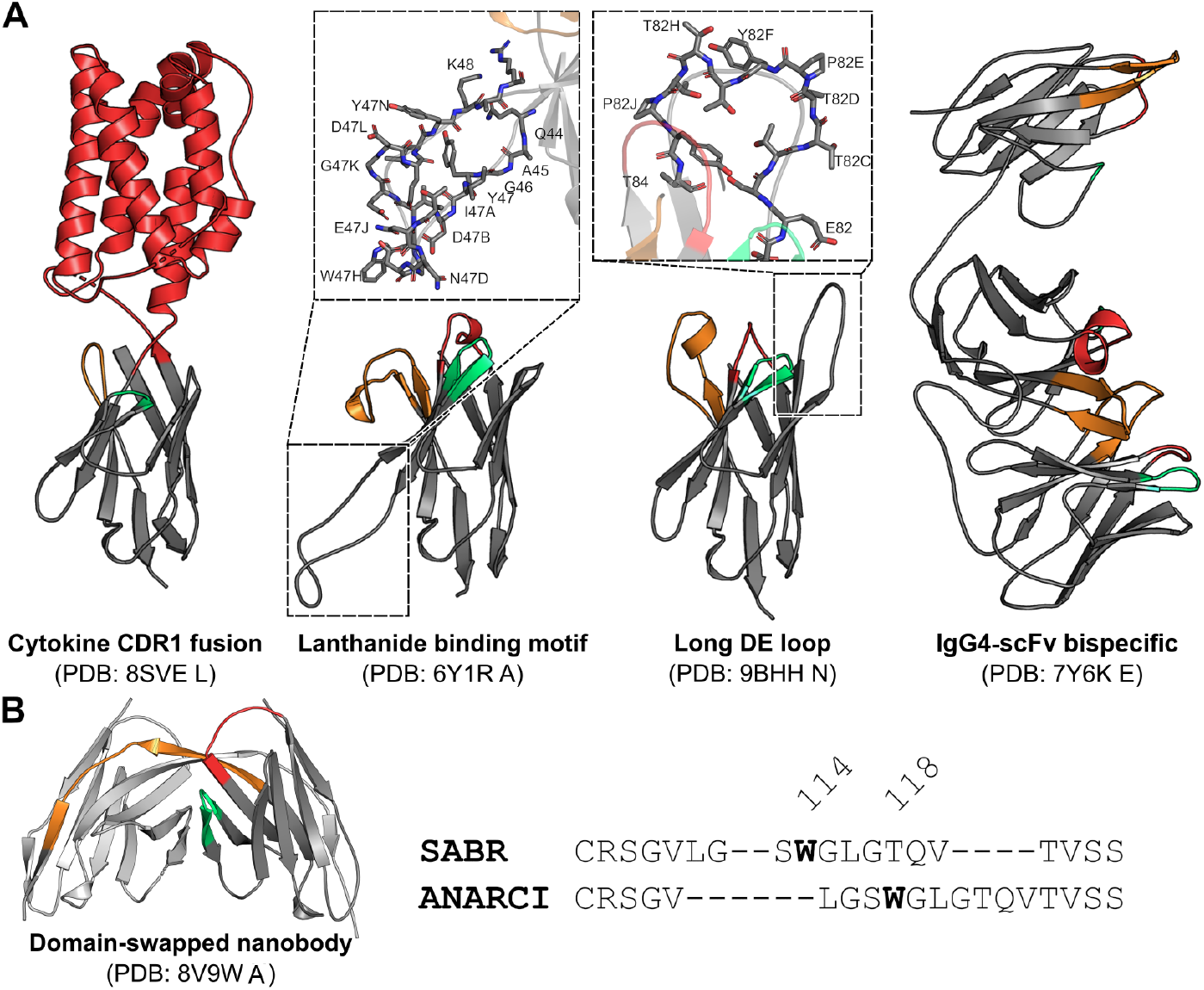
Renumbering performance on unusual one-off structures in the PDB. CDRs 1, 2, and 3 are shown in red, teal, and orange, respectively. **A)** Structures in the PDB that were correctly renumbered with SAbR despite having unusual structural features. **B)** A domain-swapped nanobody structure that SAbR misnumbered due to disruption to its canonical fold. The conserved J-gene tryptophan at IMGT position 118 is highlighted.

## 6 Conclusion

Antibody numbering can be reliably inferred from complete backbone coordinates alone, including in cases with atypical sequences or unnatural structural features. SAbR is envisioned to provide a practical annotation approach for large-scale design of engineered antibodies, particularly *de novo*–designed antibodies without obvious sequence-level clues for traditional approaches.

During the course of this work, SAbR was found to disagree with sequence-based methods on several experimental structures that had previously been flagged by a large-scale structure prediction study as having potential register misassignment errors [41]. Manual inspection corroborated those claims, providing an explanation for why sequence- and structure-based renumbering might yield divergent results without necessarily implying that either method is incorrect (Figure S4). This suggests that disagreements between SAbR and sequence-derived annotations may be used to flag candidate register errors for structural review. A larger-scale examination is left for future work.

## Acknowledgments

The author thanks Dr. Jiayi Cox and Dr. Yves Fomekong Nanfack for critical feedback on the manuscript and helpful discussions.

## A Supplementary figures

**Figure S1:**
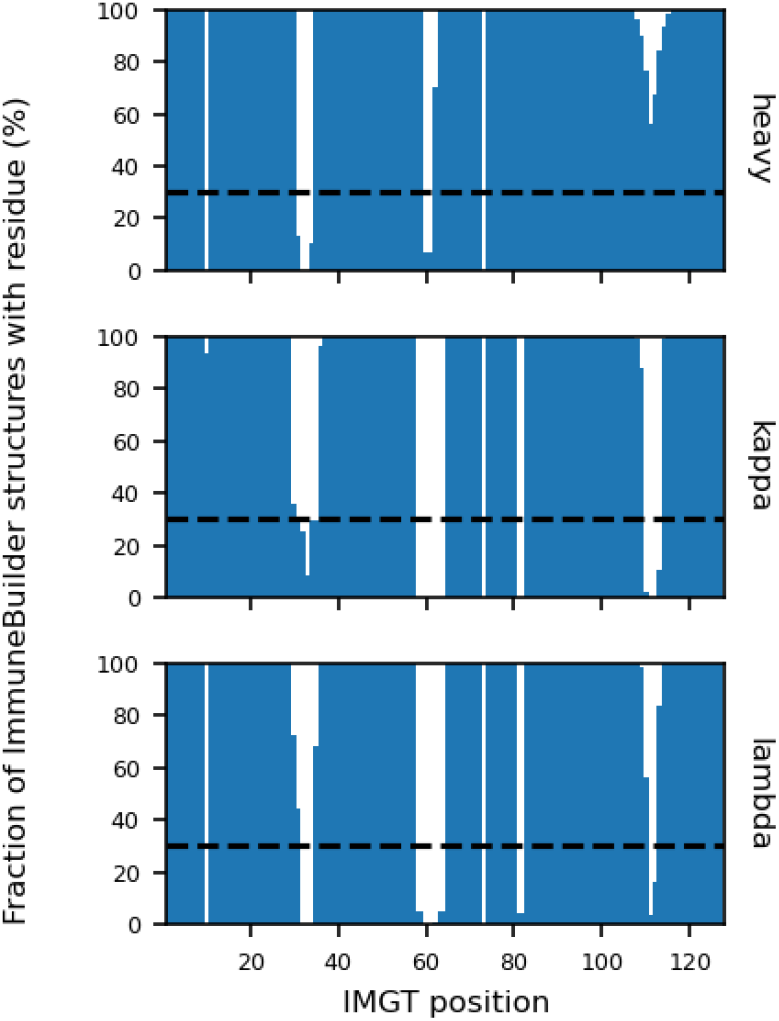
Percentage of ImmuneBuilder models across each chain type with residues mapping to specific IMGT indices.

**Figure S2:**
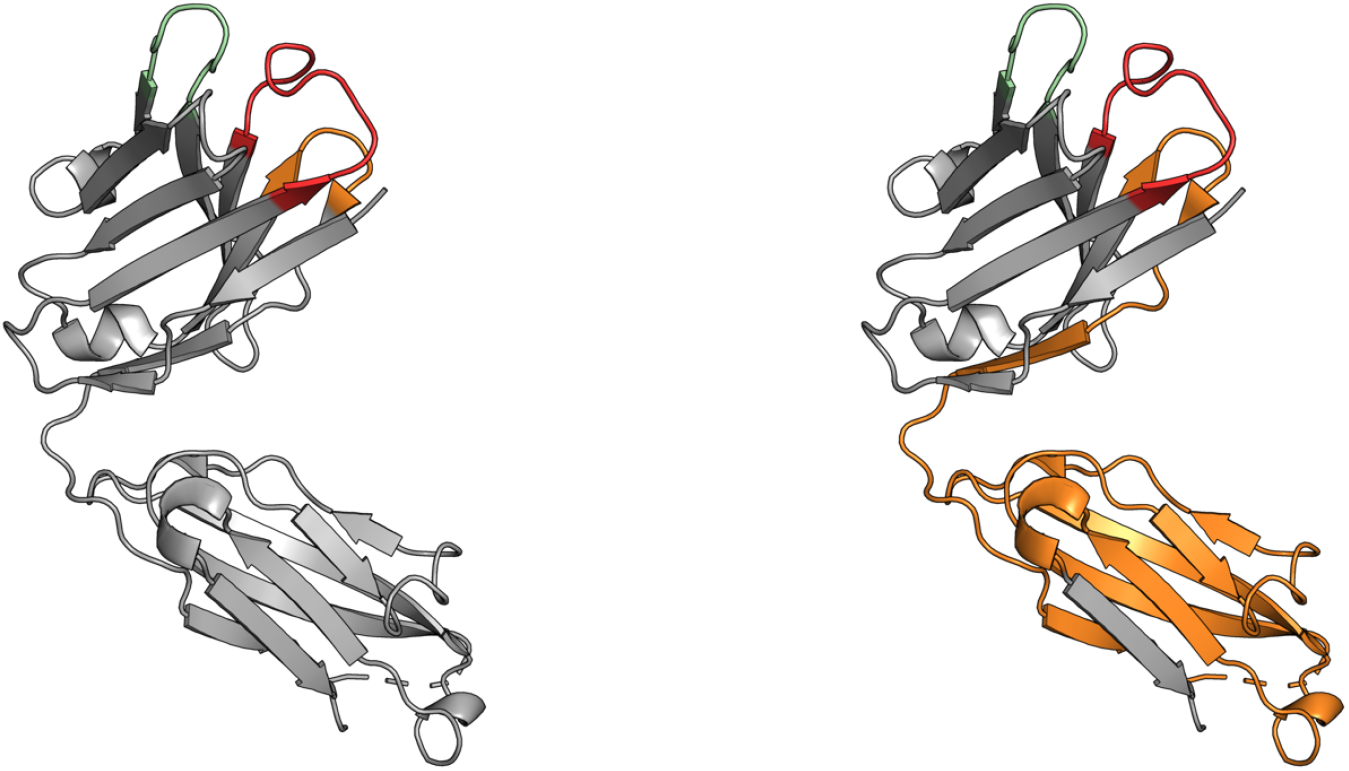
Renumbering of a Fab using default SoftAlign weights sometimes leads to assignment of huge CDR loops. **Left:** Assignment of CDR loops in PDB 7RT9 chain A in SAbDab. SAbR, when using fine-tuned weights, reproduces this assignment. Assigned CDR loops are shown in red, teal, and orange for CDRs 1, 2, and 3, respectively. **Right:** Assignment by SAbR using default SoftAlign weights. SoftAlign’s learned gap penalties include a bonus for gap extensions, leading to mistakenly assigning most of the CH1 domain to the CDR3 loop.

**Figure S3:**
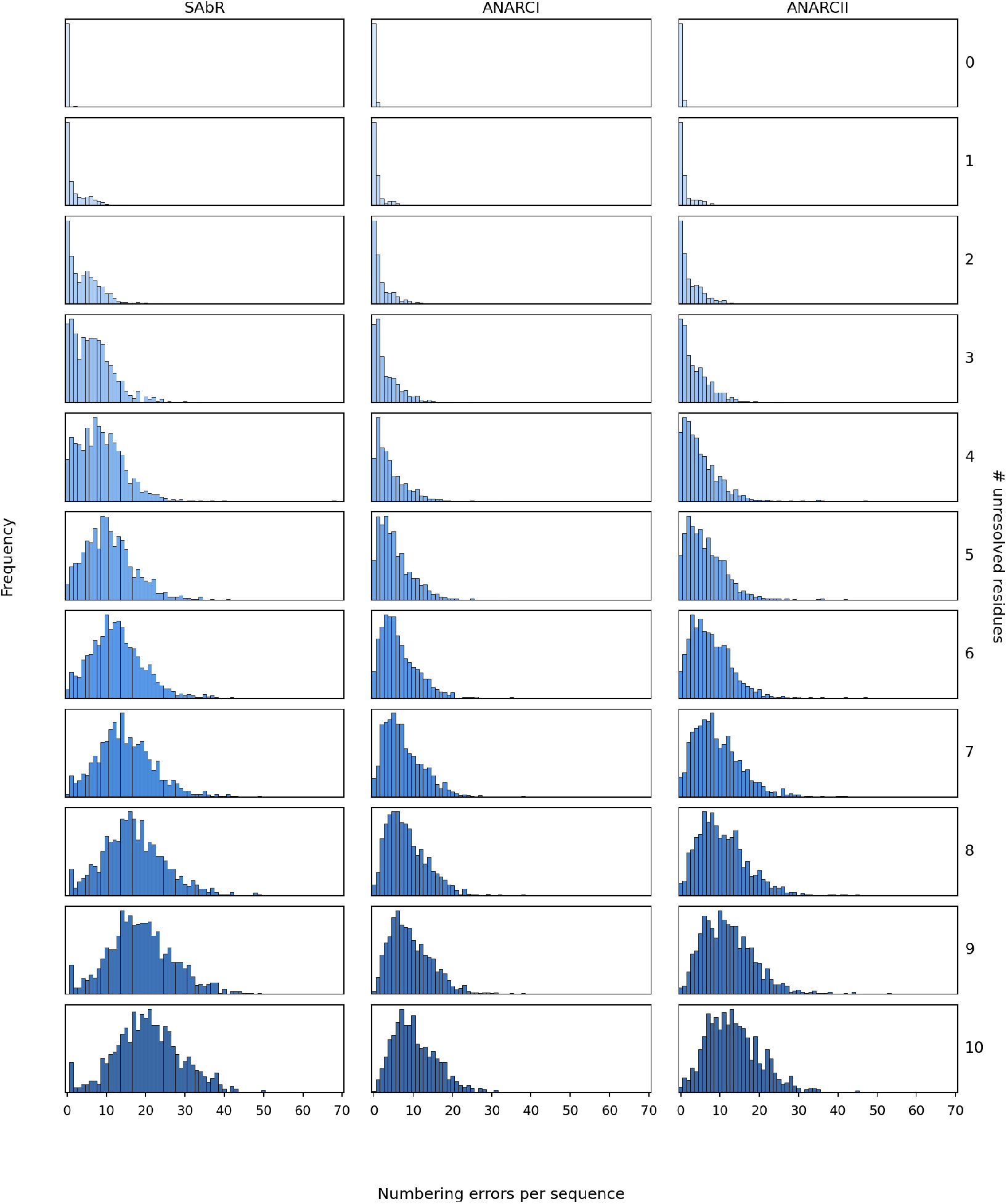
Numbering errors increase when structural gaps are present. Gaps were introduced by randomly removing residues from the variable domain of the query structure.

**Figure S4:**
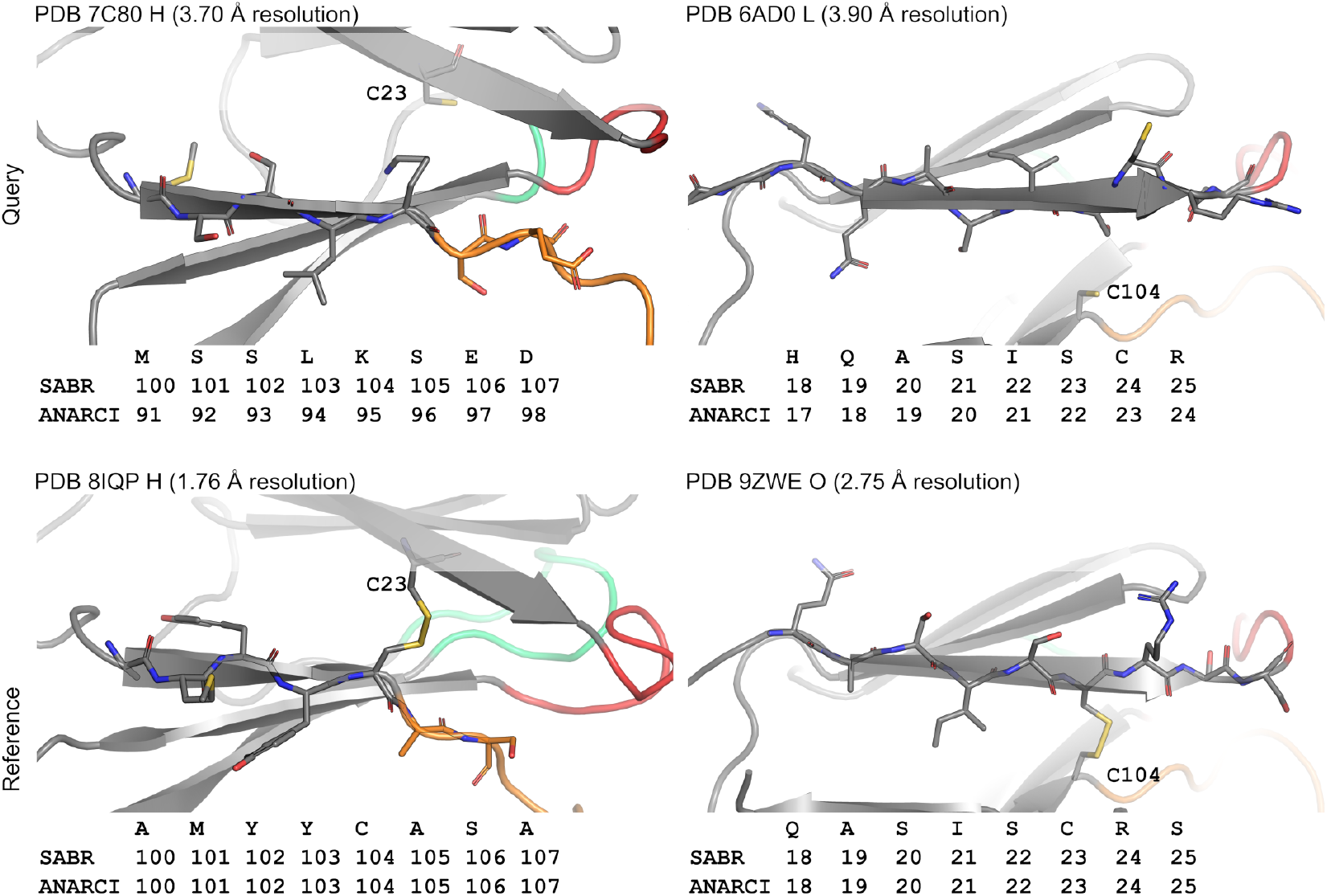
Deviations between sequence- and structure-based renumbering identify previously reported register misassignment errors in the PDB. Top and bottom panels show flagged query structures and high-resolution deviation-free reference structures, respectively. Left and right panels show two examples with deviations near the beta-strand immediately preceding the CDR3 and CDR1, respectively. A lysine and serine are placed where conserved cysteines are expected to be found. SAbR renumbers the cysteine based on its backbone position, while ANARCI renumbers it based on its identity.

## B Supplementary tables

**Table S1:** Per-residue numbering agreement outside the CDRs and DE loop across held-out and out-of-distribution test sets. The TCR evaluation comprises 73,563 residues across 946 chains and additionally excludes the C^*′′*^ strand and IMGT position 128; all structure-based TCR results use kappa-chain templates. TM-align and SoftAlign results use a single template. SAbR-SA denotes SAbR with SoftAlign weights.

|  | SAbDab<br>*n* = 1,141 | IMGT/3D<br>*n* = 42 | TCR<br>*n* = 946 | VNAR<br>*n* = 36 | Bovine<br>*n* = 37 | Real scFv<br>*n* = 57 | Synth scFv<br>*n* = 666 | *De novo*<br>*n* = 3,380 |
| --- | --- | --- | --- | --- | --- | --- | --- | --- |
| ANARCI | 99.4% | 99.1% | 99.8% | N/A | 99.9% | 100.0% | 99.9% | 85.1% |
| ANARCII | 99.8% | 99.1% | 99.6% | 97.4% | 98.1% | 100.0% | 99.9% | 71.5% |
| TM-align | 93.6% | 98.4% | 99.0% | 98.4% | 99.7% | 66.3% | 99.4% | 99.8% |
| SoftAlign | 93.8% | 98.7% | 97.4% | 98.4% | 99.8% | 75.8% | 99.5% | 99.8% |
| Foldseek | 96.1% | 98.7% | 96.9% | 96.8% | 98.8% | 84.0% | 99.2% | 98.4% |
| (3Di+AA) | 96.6% | 98.8% | 96.6% | 97.6% | 99.2% | 85.4% | 99.5% | 99.8% |
| SAbR | 99.5% | 99.1% | 98.2% | 100.0% | 99.9% | 99.5% | 99.8% | 99.9% |
| SAbR-SA | 98.4% | 99.1% | 97.2% | 97.5% | 99.8% | 98.4% | 98.8% | 100.0% |

**Table S2:** Percentage of test set structures with identical full-variable-region numbering to reference annotations. Single-template results are shown for TM-align and SoftAlign. All structure-based TCR and VNAR results use kappa-chain templates. SAbR-SA denotes SAbR with SoftAlign weights. *De novo* results include only AbMPNN-designed sequences.

|  | SAbDab<br>*n* = 1,141 | IMGT/3D<br>*n* = 42 | TCR<br>*n* = 946 | VNAR<br>*n* = 36 | Bovine<br>*n* = 37 | Real scFv<br>*n* = 57 | Synth scFv<br>*n* = 666 | *De novo*<br>*n* = 3,380 |
| --- | --- | --- | --- | --- | --- | --- | --- | --- |
| ANARCI | 98.4% | 95.2% | 96.2% | N/A | 94.6% | 96.5% | 98.5% | 40.4% |
| ANARCII | 98.2% | 95.2% | 95.9% | 94.4% | 83.8% | 96.5% | 97.6% | 39.4% |
| TM-align | 10.3% | 64.3% | 0.0% | 0.0% | 0.0% | 0.0% | 6.5% | 0.0% |
| SoftAlign | 19.5% | 69.0% | 0.0% | 0.0% | 0.0% | 0.0% | 8.7% | 0.1% |
| Foldseek | 12.8% | 69.0% | 0.0% | 2.8% | 0.0% | 1.8% | 2.4% | 0.1% |
| (3Di+AA) | 12.3% | 71.4% | 0.0% | 2.8% | 0.0% | 0.0% | 2.9% | 0.1% |
| SAbR | 96.2% | 95.2% | 0.0% | 0.0% | 94.6% | 89.5% | 93.8% | 99.4% |
| SAbR-SA | 90.9% | 95.2% | 0.0% | 0.0% | 81.1% | 71.9% | 70.4% | 99.9% |

**Table S3:**
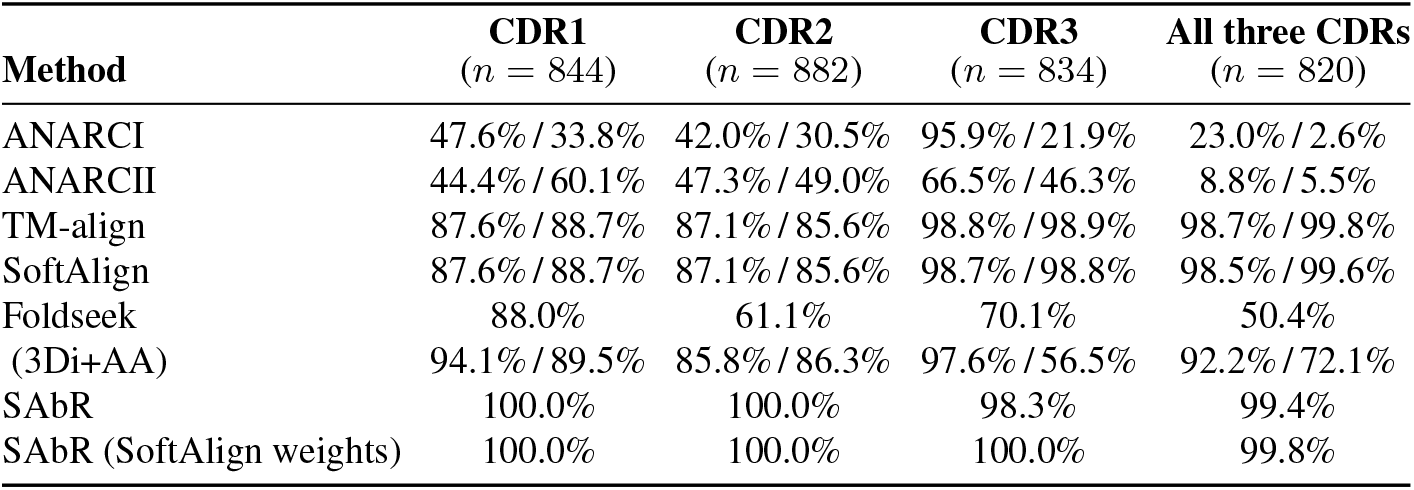
Percentage of expected RFantibody designs with zero framework numbering errors. Paired values are reported as AbMPNN/poly-Gly; single values are reported where sequence variants are not separated. Missing predictions, numbering failures, and missing sequence-variant counterparts are counted as failures, so paired values use common denominators. TM-align and SoftAlign use a single template selected separately for each sequence variant.

| Method | CDR1<br>(*n* = 844) | CDR2<br>(*n* = 882) | CDR3<br>(*n* = 834) | All three CDRs<br>(*n* = 820) |
| --- | --- | --- | --- | --- |
| ANARCI | 47.6% / 33.8% | 42.0% / 30.5% | 95.9% / 21.9% | 23.0% / 2.6% |
| ANARCII | 44.4% / 60.1% | 47.3% / 49.0% | 66.5% / 46.3% | 8.8% / 5.5% |
| TM-align | 87.6% / 88.7% | 87.1% / 85.6% | 98.8% / 98.9% | 98.7% / 99.8% |
| SoftAlign | 87.6% / 88.7% | 87.1% / 85.6% | 98.7% / 98.8% | 98.5% / 99.6% |
| Foldseek | 88.0% | 61.1% | 70.1% | 50.4% |
| (3Di+AA) | 94.1% / 89.5% | 85.8% / 86.3% | 97.6% / 56.5% | 92.2% / 72.1% |
| SAbR | 100.0% | 100.0% | 98.3% | 99.4% |
| SAbR (SoftAlign weights) | 100.0% | 100.0% | 100.0% | 99.8% |

**Table S4:**
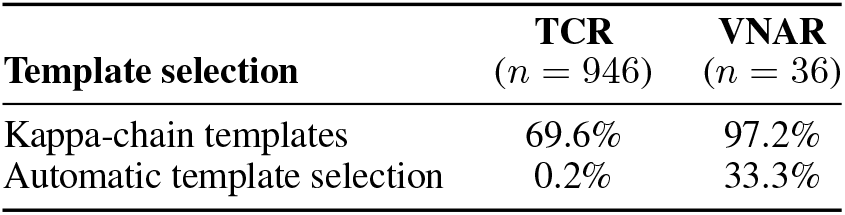
Effect of template selection on SAbR performance for TCRs and shark VNARs. Values are the percentage of structures with zero framework numbering errors.

| Template selection | TCR<br>(*n* = 946) | VNAR<br>(*n* = 36) |
| --- | --- | --- |
| Kappa-chain templates | 69.6% | 97.2% |
| Automatic template selection | 0.2% | 33.3% |

**Table S5:**
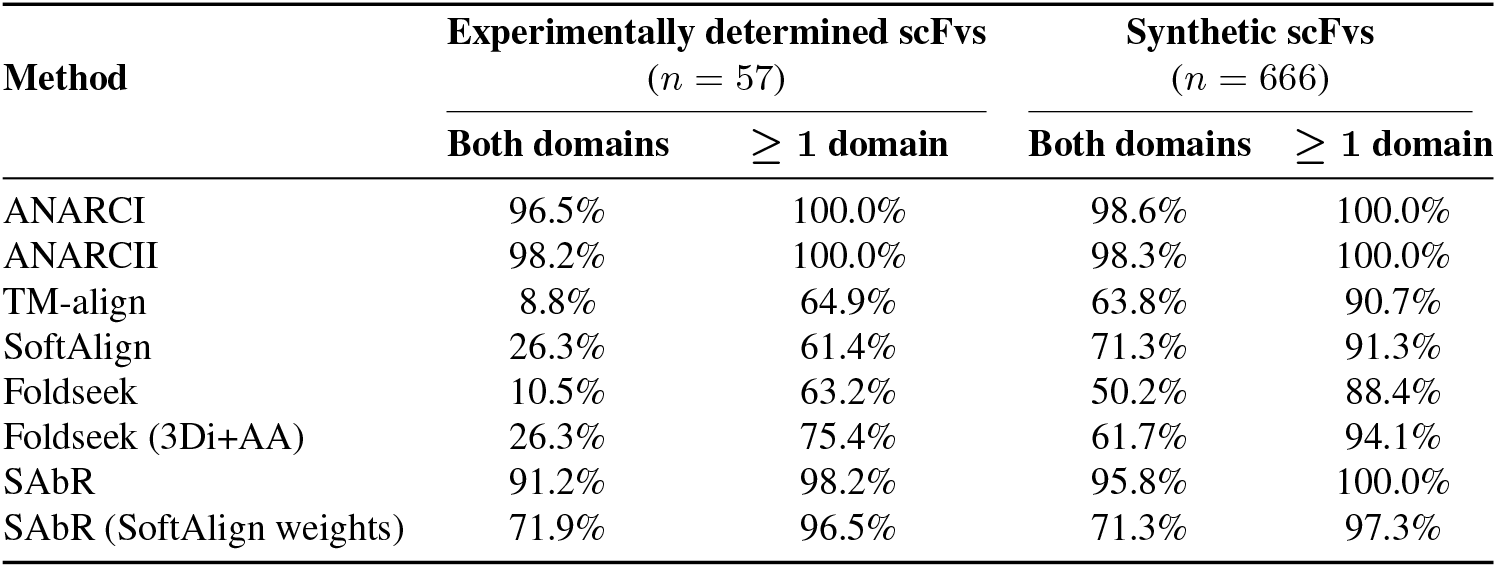
Percentage of experimentally determined and synthetic scFvs with zero framework numbering errors in both variable domains or in at least one domain. TM-align and SoftAlign results use a single template.

**Table S6:**
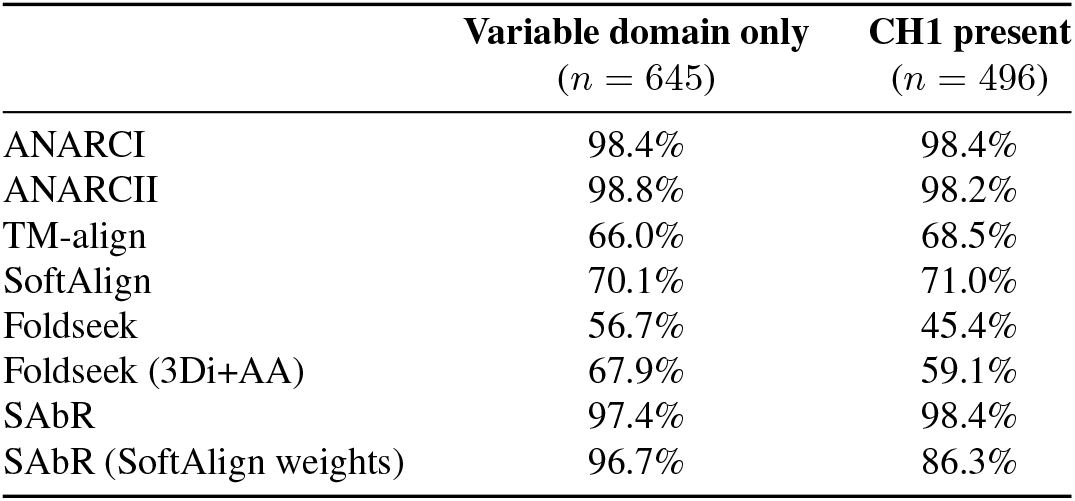
Percentage of SAbDab test structures with zero framework numbering errors, stratified by the presence of the C-terminal CH1 domain. TM-align and SoftAlign results use a single template. SAbR-SA denotes SAbR with SoftAlign weights. ANARCI was used to number the SAbDab reference sequences.

